# The hippocampus and neocortical inhibitory engrams protect against memory interference

**DOI:** 10.1101/366377

**Authors:** R.S. Koolschijn, U.E. Emir, A.C. Pantelides, H. Nili, T.E.J. Behrens, H.C. Barron

**Author notes:** indicates equal contribution (T.E.J. Behrens, H.C. Barron). Corresponding authors: Renée S. Koolschijn and Helen C. Barron.

## Abstract

Our experiences often overlap with each other, sharing features, stimuli or higher-order information. But despite this overlap, we are able to selectively recall individual memories to guide our decisions and future actions. The neural mechanisms that support such precise memory recall, however, remain unclear. Here, using ultra-high field 7T MRI we reveal two distinct mechanisms that protect memories from interference. The first mechanism involves the hippocampus, where the BOLD signal predicts behavioural measures of memory interference, and contextual representations that aid separation of overlapping memories are organised using a relational code. The second mechanism involves neocortical inhibition: when we reduce the concentration of neocortical GABA using trans-cranial direct current stimulation (tDCS) neocortical memory interference increases in proportion to the reduction in GABA, which in turn predicts behavioural performance. Together these findings suggest that memory interference is mediated by both the hippocampus and neocortex, where the hippocampus aids separation of memories by coding context-dependent relational information, while neocortical inhibition prevents unwanted co-activation between overlapping memories.

## Introduction

Our decisions and actions are often guided by past experiences that overlap with one another in their content or sensory information. To ensure that interference between related or overlapping experiences is minimised, a stable memory storage system is critical. However, the precise physiological mechanism that supports stable memory storage in the absence of memory interference remains unclear.

One way to minimize memory interference is to separate stored information using contextual representations (McClelland et al., 1995; Norman and O’Reilly, 2003; Shapiro and Olton, 1994). Behavioural data in humans provides supporting evidence for this mechanism, as contextual cues help mitigate memory interference between two lists of paired associates (Bilodeau and Schlosberg, 1951). At the neural level, anticorrelated firing patterns for opposing contexts can be observed in the hippocampal output regions (Butterly et al., 2012; McKenzie et al., 2014). These contextual representations are thought to emerge from competitive pattern separation of overlapping memories (Yassa and Stark, 2011). However, this contrasts with an alternative view of the hippocampus which suggests that overlapping memories are integrated into a common relational framework (Eichenbaum, 2004). To reconcile these two possibilities, here we hypothesise that the hippocampus helps protect against memory interference by binding memories to contextual representations that are then organised according to the relational structure of learned information.

In addition, an alternative way to protect stored memories from interference involves using inhibition. Following learning, new information is thought to be stored in the brain via modification in the strength of excitatory connections (Hebb, 1949; Nabavi et al., 2014; Song and Abbott, 2001). In turn, these newly modified excitatory connections are opposed by equivalent changes in the strength of inhibitory connections (Barron et al., 2016a; Froemke et al., 2007; Vallentin et al., 2016; Vogels et al., 2011). This allows excitatory-inhibitory (EI) balance to be maintained despite new learning (Froemke et al., 2007; Haider et al., 2006; Okun and Lampl, 2008; Wehr and Zador, 2003), and ensures that memories lie dormant unless EI balance is disturbed (Barron et al., 2016a; Jacobs and Donoghue, 1991; Vallentin et al., 2016). Here we hypothesize that the inhibitory component of a memory, otherwise termed the inhibitory engram (Barron et al., 2017), protects memories from interference by preventing run-away excitation.

Consistent with this hypothesis, context-dependent behaviour in rodents is accompanied by modulation of neocortical interneurons (Kuchibhotla et al., 2017), while in humans an increase in neocortical GABA relative to glutamate accompanies overlearning, a process known to protect memories from interference (Shibata et al., 2017). Clinical investigations also support a key role for inhibitory regulation of memory expression, as impaired GABAergic regulation can readily account for delusions and hallucinations reported in schizophrenia (Vogels and Abbott, 2007; Yizhar et al., 2011). Thus, by gating memory expression (Barron et al., 2016a, 2017; Vogels and Abbott, 2009), inhibitory engrams may play a critical role in preventing unwanted interference between overlapping memories.

Here we investigate the role of both the hippocampus and neocortical inhibition in protecting against memory interference. First, we test the hypothesis that contextual representations in the hippocampus are organised using a relational code, thus separating competing memories according to behaviourally relevant information. Second, we test the hypothesis that neocortical inhibition protects overlapping memories from interference.

To this end, we designed a task that required participants to encode two overlapping but context-dependent memories across two consecutive days. On the third day, interference between the two memories was measured using ultra-high field 7T Magnetic Resonance Imaging (MRI). In the hippocampus, we observed an increase in hippocampal Blood-Oxygen-Level Dependent (BOLD) signal during opportunities for memory interference which predicted subsequent behavioural performance. In addition, the underlying contextual representations were organised according to the relational structure of the two memories. To then investigate the role of neocortical inhibition in protecting memories from interference, half way through the scan we manipulated the concentration of neocortical GABA using brain stimulation, and re-assessed evidence for memory interference. The drop in neocortical GABA induced by brain stimulation predicted an increase in neocortical memory interference, which in turn predicted deficits in behavioural performance. Together these results suggest that memory interference is mediated by two distinct mechanisms: a hippocampal mechanism where contextual representations are organised according to behaviourally relevant relationships, and a neocortical mechanism where inhibition protects overlapping memories from unwanted co-activation.

## Results

### Associative learning and experimental design

On day 1 of the experiment, participants learned a set of associations between seven rotationally-invariant abstract stimuli (Fig. 1A) which together formed ‘memory 1’. Within memory 1 each stimulus was associated with two other stimuli, giving seven bidirectional associations in total. The set of associations could be arranged into a ring structure (Fig. 1B), although this was never made explicit to the participants. Instead, participants were instructed to learn the associations using a three-alternative forced choice task (Fig. 1D, Supplementary Fig. 2A, see Methods).

**Figure 1:**
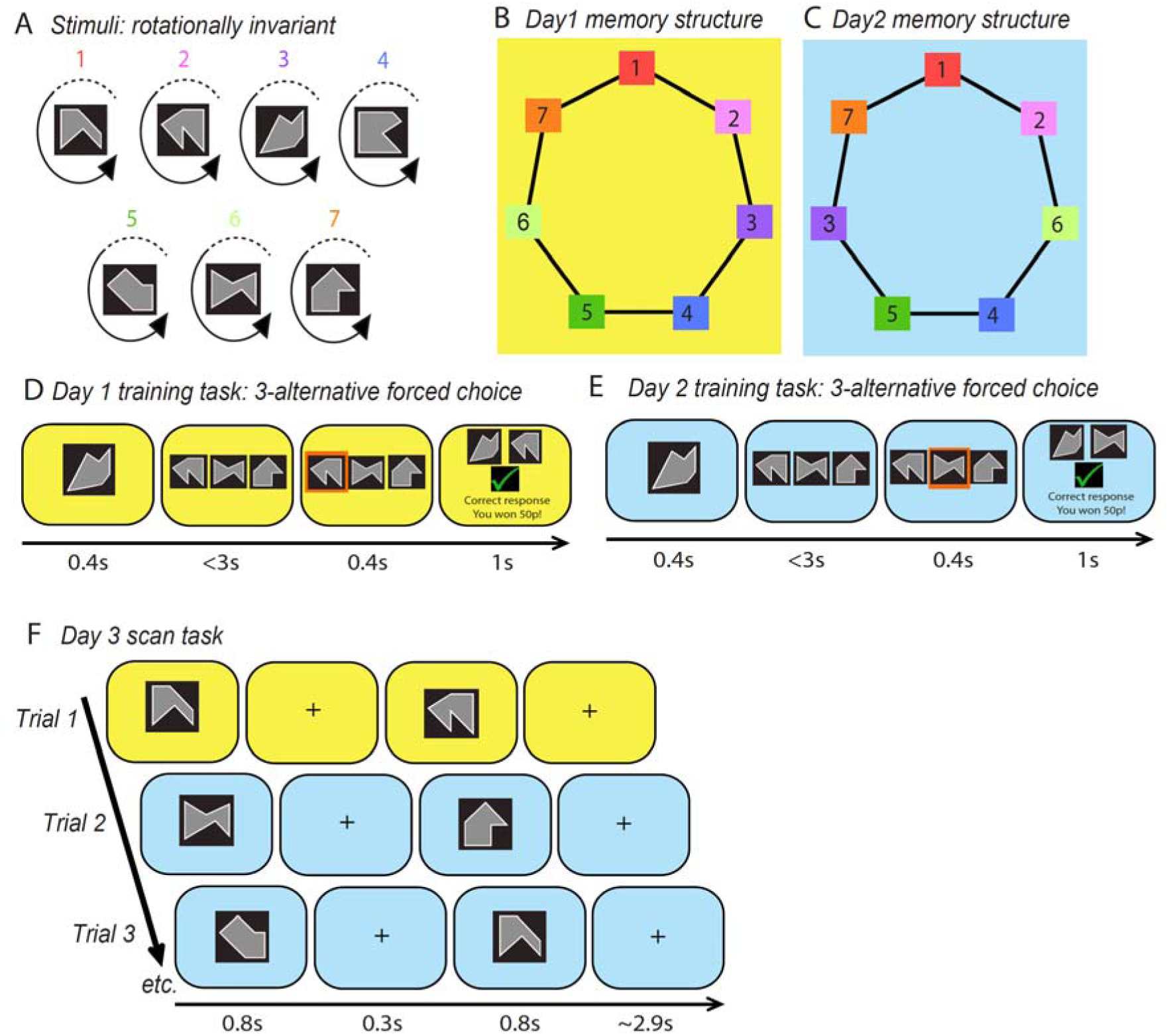
Experimental design. **A**) Seven stimuli were used for the experiment, where each stimulus was an abstract shape which could appear in four possible rotations. **B-E**) Example experimental protocol; **B**) On day 1, participants learned seven associations between pairs of the seven stimuli shown in *A*, and a contextual cue was provided using a yellow background. The associations could be arranged in a ring structure, although this was not explicitly shown to participants. **C**) On day 2, participants learned seven associations between pairs of the seven stimuli, four of which were different from the associations learned on day 1 as the position of stimuli 3 and 6 were swapped. A contextual cue was provided using a blue background, a different colour from that used on day 1. **D-E**) To learn the associations on day 1 (D) and day 2 (E), participants performed a 3-alternative forced choice task where the appropriate background colour was used to provide a contextual cue. **F**) During the scan task, participants observed pairs of stimuli presented consecutively against either a yellow or blue background. All possible pairs of stimuli were presented in a random order.

On day 2 of the experiment, participants learned a second set of bidirectional associations between the same seven abstract stimuli (Fig. 1A) which together formed ‘memory 2’. As in memory 1, each stimulus in memory 2 was associated with two other stimuli (Fig. 1C). Participants again learned these associations using the three-alternative forced choice task (Fig. 1E, Supplementary Fig. 2B). Critically, the relative position of the seven stimuli within the ring differed between memory 1 and memory 2, as the position of stimuli three and six was switched (Fig. 1B-C). Consequently, four of the seven associations in memory 2 were different from those in memory 1, while three associations remained the same. To help participants distinguish between memory 1 and memory 2, contextual cues were used, consisting of a unique background colour (yellow or blue, randomised across participants) (Fig. 1D-E) and a time interval of approximately 24 hours between learning sessions.

Thus, memory 1 and memory 2 included the same stimuli but had different relational structures. The difference in relational structure was designed to ensure that a subset of associations across memory 1 and 2, those containing stimuli three or six, were different, while the remaining associations were matched. We predicted that associations containing elements three or six were susceptible to memory interference, where memory interference manifests as recall of a relational neighbour from the alternative, inappropriate memory. Meanwhile the matched portion of the two memories provided the necessary control. The experimental design therefore included precise and controlled markers of memory interference that could be assessed at both a behavioural and neural level.

### Hippocampus mediates memory interference using context-dependent relational codes

To identify the physiological mechanisms that protect memories from interference, we first considered the contribution made by the hippocampus. One possibility is that the hippocampus helps prevent memory interference by binding memories to contextual representations that reflect the output of competitive pattern separation (Yassa and Stark, 2011). While opposing contextual representations have been identified in distinct ensembles of cells in the hippocampus (McKenzie et al., 2014), an alternative view of the hippocampus suggests that overlapping memories are integrated into a common relational code (Eichenbaum, 2004). Here, in an attempt to reconcile these two possibilities, we predicted that the hippocampus binds memories to contextual representations which are then organised according to behaviourally relevant relationships.

To test this prediction, on day 3 of the experiment we used fMRI to measure the BOLD response to the associative memories learned in memory 1 and 2. On each trial of the scan task, a pair of stimuli was presented on either a yellow or a blue background to provide a contextual cue for memory 1 or memory 2 (Fig. 1F, see Methods). We controlled for potential confounds introduced by expectation suppression (Summerfield et al., 2008) by ensuring that each possible pair of stimuli was presented equally often in a fully randomized order. To ensure participants paid close attention to the stimuli presented during the scan, participants were instructed to detect ‘odd-ball’ stimuli which were not part of the seven experienced during training.

We observed an increase in the hippocampal BOLD signal to pairs of stimuli that had a different relative position across memory 1 and 2 (i.e. pairs of stimuli that included stimuli 3 and 6), relative to pairs of stimuli that had the same relative position across both memories (right hippocampus: p=0.015, FWE corrected, peak t_23_=4.34, Fig. 2A; left hippocampus: p=0.056 FWE corrected, peak t_23_=3.66, Fig. 2B; corrected for multiple comparisons using small volume correction, Fig. 2D, see Methods; both hippocampi: Fig. 2C, for visualisation). Therefore, the hippocampal BOLD signal increased with opportunity for memory interference.

**Figure 2:**
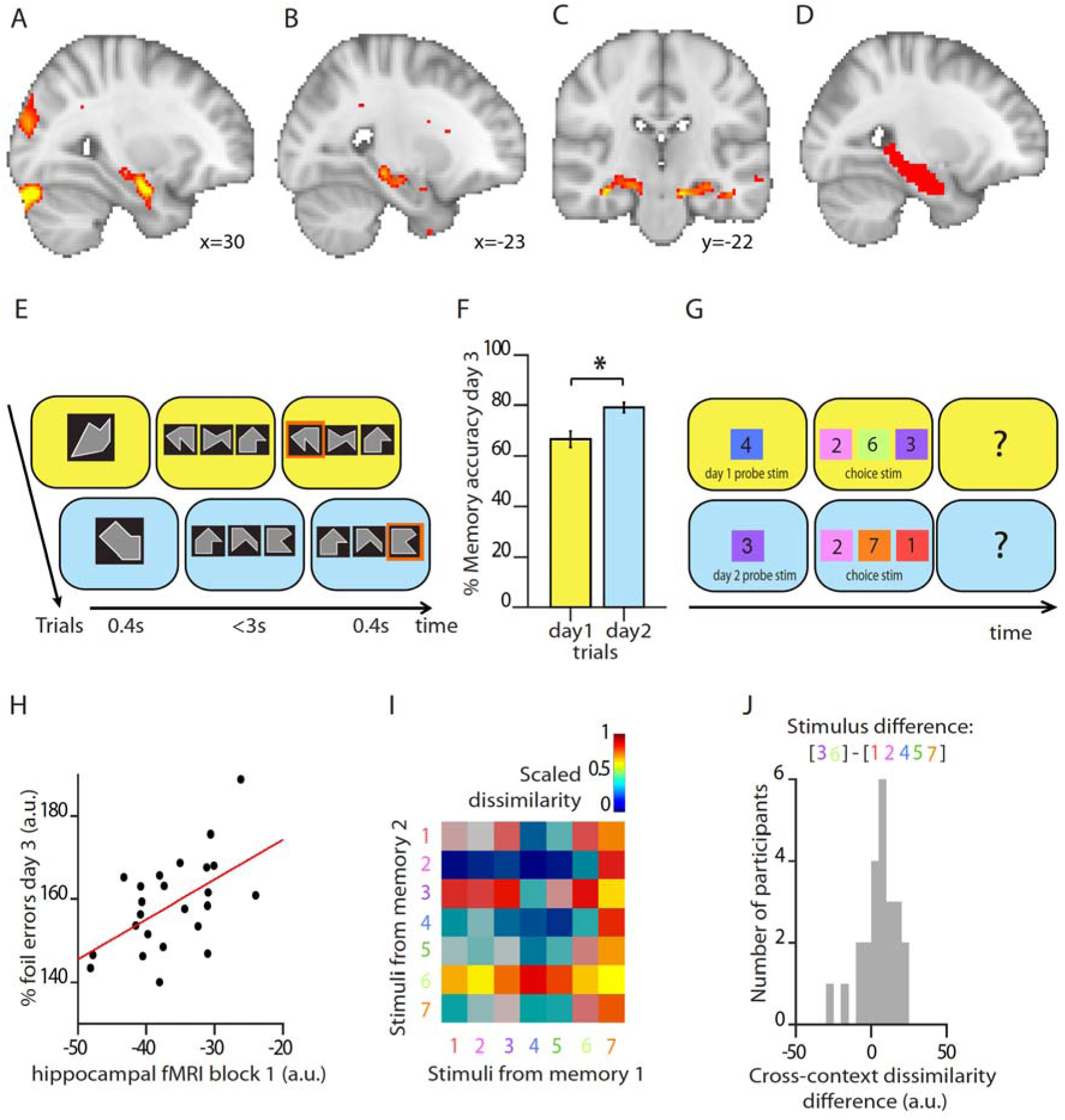
The hippocampus mediates memory interference by representing context-dependent relational information. **A-C**) Hippocampal BOLD signal was higher on trials where there was opportunity for memory interference, i.e. when trials include stimuli 3 or 6 that have a different relative position between memory 1 and 2 (thresholded at p<0.01 for visualization). **A-B**) Hippocampal BOLD signal was significantly higher in right hippocampus (p=0.015, FWE corrected, peak t_23_=4.34, *A*), while a similar trend was observed in the left hippocampus (p=0.056, FWE corrected, peak t_23_=3.66, *B*). **C**) Hippocampal BOLD signal across both hippocampi, for visualisation. **D**) Hippocampal ROI used to perform SVC method in *A-B*, and to extract parameter estimates in *H*. **E**) After exiting the scanner on day 3, participants performed a surprise memory test. On each trial participants were presented with a probe stimulus with the background colour providing the contextual cue. They were then presented with three option stimuli and were required to choose the stimulus correctly paired with the probe stimulus in the absence of feedback. **F**) Memory accuracy on the surprise memory test (mean, ± SEM), for memory 1 (day 1) and memory 2 (day 2). The more recent associations in memory 2 were remembered more accurately than those in memory 1 (paired t-test: t_2_s=3.99, p<0.001). **G**) Foil trials on the surprise memory test shown in *E* were identified as those trials where one of the three stimuli was incorrect given the current context, but correct in the alternative context. **H**) The hippocampal BOLD response to trials where there was opportunity for memory interference during the first scan predicted behavioural memory interference on the post-scan surprise memory test (Pearson’s correlation: r_23_=0.54, p=0.006). **I**) Representational Similarity Analysis was used to extract the representational dissimilarity between memory 1 and 2 for each of the seven stimuli. For each stimulus representation, all trials containing the stimulus were included, e.g. stimulus 1 in memory 1 included all pairs of stimuli shown on a yellow background that included stimulus 1, i.e. 1-1, 1-2, … 1-7. The confusion matrix shown here illustrates the cross-memory representational dissimilarity for all seven stimuli, averaged across all participants, rank transformed and scaled into [0-1] for visualisation. **J**) Histogram across all participants showing the difference in hippocampal representational dissimilarity between memory 1 and 2 for stimuli that had different relative position across memory 1 and 2 (stimuli 3 and 6) versus those stimuli that had the same relative position across memory 1 and 2 (stimuli 1,2,4,5,7) (stimuli 3/6 - stimuli 1/2/4/5/7: Wilcoxon sign rank test: Z_23_=2.2, p=0.028).

We next asked whether this hippocampal BOLD signal could predict behavioural measures of memory interference. After the scan session, participants performed a surprise memory test and we assessed recall accuracy for all seven associations within memory 1 and memory 2. The memory test involved the three-alternative force choice task used during training, but now in the absence of feedback (Fig. 2E). On average, participants correctly recalled the appropriate association on 74.3% of trials, showing higher accuracy for more recent memories (paired-sample t-test, t_25_=3.99, p<0.001, Fig. 2F). In addition to participants’ overall memory accuracy, behavioural memory interference was quantified using participants’ performance on ‘foil trials’, namely those trials where the choice stimuli included the stimulus that was correct given the contextual background cue, but also a ‘foil’ stimulus that would be correct in the alternative memory (Fig. 2G). The percentage of ‘foil errors’ made by a participant corresponded to the percentage of foil trials where the foil stimulus was chosen rather than the correct stimulus. Across participants, we observed a positive relationship between the hippocampal BOLD signal (Fig. 2A-C) and this behavioural index for memory interference (hippocampal BOLD Block 1 vs. foil errors: r_23_=0.54, p=0.006, Fig. 2H, after accounting for differences in learning).

While these results suggest that the hippocampus may play a key role in mediating memory interference, they leave open the nature of the hippocampal code. If the hippocampus embeds memories in contextual representations that are organized using a relational code, then representations of stimuli that have different relative positions across memory 1 and 2 (i.e. stimuli 3 or 6) should have less similar activity patterns across the two contexts, compared to representations of stimuli that have the same relative position across memory 1 and 2 (i.e. stimuli 1, 2, 4, 5, 7).

To test this hypothesis we used representational similarity analysis (RSA) to extract the pattern of activity across voxels in a hippocampal ROI for each trial (for ROI see Fig. 2D). We then quantified the representational dissimilarity between memory 1 and memory 2 for each of the seven stimuli using the Mahalanobis distance (Fig. 2I, see Methods). Notably, for this stimulus-wise cross-memory comparison the only difference in visual information between trials in memory 1 and 2 was the background colour. Across participants we observed higher cross-memory dissimilarity for stimuli that have a different relative position across memory 1 and 2 (stimuli 3 and 6) relative to stimuli that have the same relative position across memory 1 and 2 (stimuli 1,2,4,5,7) (Fig. 2J; Wilcoxon sign rank test across the group: Z_23_=2.2, p=0.028). Furthermore, stimuli that have a different relative position across memory 1 and 2 (i.e. stimuli 3 and 6) also showed higher cross-memory dissimilarity with representations of all other possible stimuli (Supplementary Fig. 1A). This effect was particularly evident for stimuli 3 and 6 in memory 2 (rows 3 and 6 of Fig. 2I; Supplementary Fig. 1B). This suggests that the hippocampus embeds memories in contextual representations that are organized according to a relational code.

### Manipulating neocortical EI balance to measure the effect of inhibition on memory

Having characterised a role for the hippocampus in mediating memory interference, we next asked whether inhibition in the neocortex also plays a key role. Unlike the hippocampus, which appears to encode associated stimuli using conjunctive representations, associative memories in neocortex appear to be stored by excitatory connections that are later balanced by matched inhibition (Barron et al., 2016a; Froemke et al., 2007; Vallentin et al., 2016; Vogels and Abbott, 2009). Therefore, by day 3 of the experiment we expected neocortical representations of memory 1 and 2 to be stored in balanced EI ensembles. However, if neocortical inhibition plays a critical role in protecting overlapping memories from unwanted interference then it should be possible to induce interference by reducing inhibitory tone. To test this prediction, in the second half of the day 3 scan session we applied non-invasive anodal transcranial direct current stimulation (tDCS) (Fig. 3A-B), a tool previously used to induce a transient reduction in the concentration of neocortical GABA (Barron et al., 2016a; Kim et al., 2014; Stagg et al., 2009) and to unmask otherwise silent neocortical associative memories (Barron et al., 2016a).

**Figure 3:**
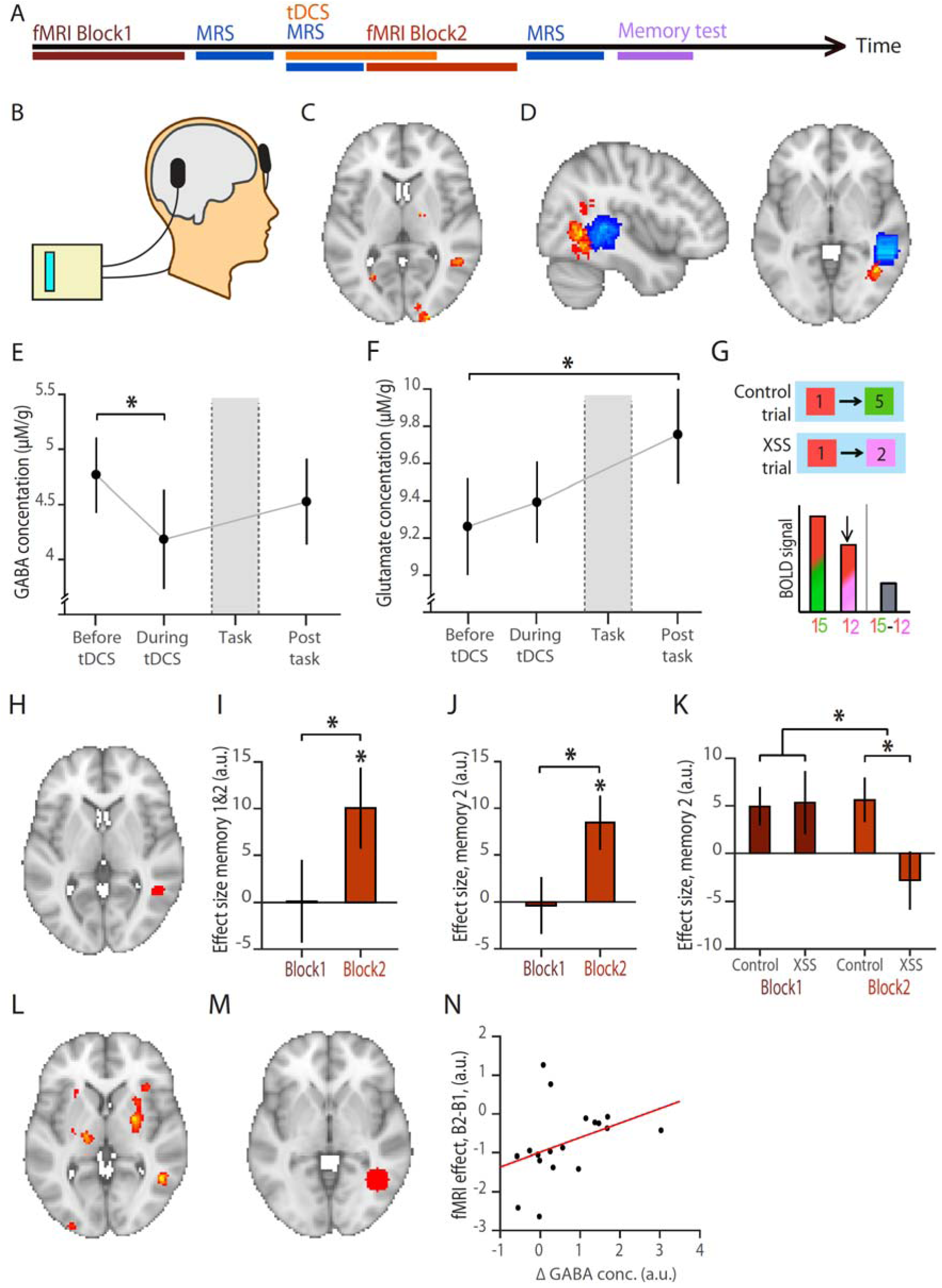
Manipulating neocortical EI balance using brain stimulation unmasks otherwise dormant neocortical memories. **A**) Schematic showing protocol used for day 3 of the experiment. Participants started with Block 1 of the scan task, as shown in Fig. 1F, before MRS measurements were taken to estimate baseline measures of 19 different metabolites. Anodal tDCS was then applied for a total of 20 minutes to induce EI imbalance, with MRS measurements taken during the first 10 minutes before Block 2 of the scan task. After Block 2 of the scan task, a third set of MRS measurements were obtained before participants exited the scanner to perform a surprise memory test. **B)** After the first scan task (Block 1), and while participants lay in the scanner, anodal tDCS was applied to the aLOC, with the cathodal electrode placed over the contralateral supraorbital ridge. **C)** Previously published data(Barron et al., 2016a) showing re-expression of associative memories during application of anodal tDCS. This aLOC region was the target location for the tDCS electrode in the current experimental protocol. **D)** Blue: Average location of the MRS voxel across participants. Red: Average location of the tDCS electrode across participants, projected into the brain (see Methods). **E)** MRS was used to quantify the concentration of GABA at three time points indicated in *A* (shown: mean ±SEM). A significant reduction in GABA was observed during tDCS (‘Before tDCS’ – ‘During tDCS’, t_19_= 1.97, p=0.016). **F)** MRS was used to quantify the concentration of glutamate at three time points indicated in *A* (shown: mean ±SEM). A significant increase in glutamate was observed after the second scan-task (‘Post-task’ – ‘Before tDCS’, t_19_= 2.26, p=0.018). **G)** When participants performed the scan task in EI imbalance, we predicted an increase in cross-stimulus suppression (‘XSS’) on trials where pairs of directly associated stimuli were presented (e.g. stimuli 1 and 2), relative to trials where pairs of unassociated stimuli were presented (‘Control’, e.g. stimuli 1 and 5). The difference between ‘Control’ and ‘XSS’ trials could be indexed using the BOLD signal to provide a measure of cross-stimulus suppression. **H)** To test replication of our previously published result(Barron et al., 2016a) (shown in **C)**, an ROI was defined by thresholding C at p<0.01 (see Methods). **I)** For directly associated stimuli across both memory 1 and 2, extracted parameter estimates (shown: mean ±SEM) revealed a significant increase in fMRI cross-stimulus suppression from Block 1 to 2, and significant fMRI cross-stimulus suppression on Block 2 (within the ROI shown in *H*: ‘Control’ – ‘XSS’ for Block 2 – Block 1: t_23_=1.73, p=0.049; for Block 2: t_23_=2.31, p=0.015). **J)** Between directly associated stimuli in memory 2, extracted parameter estimates (shown: mean ±SEM) revealed a significant increase in fMRI cross-stimulus suppression from Block 1 to 2, and significant fMRI cross-stimulus suppression on Block 2 (within the ROI shown in *H:* ‘Control’ – ‘XSS’ for Block 2 – Block 1: t_23_=2.31, p=0.016; Block 2: t_23_=2.91, p=0.004). This was not observed for memory 1 (Supplementary Fig. 4A, p>0.05). **K)** Extracted parameter estimates from *E* split into the ‘Control’ and ‘XSS’ conditions, as described in *G* (shown: mean ±SEM). **L)** T-statistic map for cross-stimulus suppression between directly associated stimuli in memory 2 during Block 2, thresholded at p<0.01 uncorrected for visualization. **M)** 10mm radius sphere defined around the peak tDCS electrode location for all participants (see Methods). **N)** There was a significant positive correlation between the change in GABA (‘Before tDCS’ – ‘During tDCS’) and the increase in fMRI cross-stimulus suppression (Block 2-Block 1) observed in the ROI shown in *M*, averaged across both memory 1 and 2 *(B* indicates ‘Block’, Spearman correlation: r_17_=0.52, p=0.028, after accounting for changes in glutamate, see Methods).

Direct current stimulation increases cortical excitability, such that neuronal firing rates increase (Bindman et al., 1962) along with remote motor evoked potentials measured using transcranial magnetic stimulation (TMS) (Nitsche et al., 2005). After stimulation, the increase in cortical excitability is sustained for minutes to hours (Bindman et al., 1962) via a protein synthesis dependent process (Nitsche and Paulus, 2000), which can be used to enhance learning (Jacobson et al., 2011) and recovery from stroke (Hummel and Cohen, 2006). Critically, the mechanism responsible for this increase in cortical excitability appears to involve a reduction in the available GABA concentration, as evidenced by in vivo spectroscopic measurements (Barron et al., 2016a; Kim et al., 2014; Stagg and Nitsche, 2011; Stagg et al., 2009).

Taking advantage of this non-invasive tool here, we placed the anodal tDCS electrode over anterior Lateral Occipital Complex (aLOC) to target the brain region known to encode associative memories between rotationally invariant shapes (Barron et al., 2016a) (Fig. 3C). The cathodal electrode was placed over the contralateral supraorbital ridge (Fig. 3B). Brain stimulation was applied immediately before participants performed a second run of the scan task (block 2, Fig. 3A). Before, during and after brain stimulation (Fig. 3A) we used Magnetic Resonance spectroscopy (MRS) to rapidly measure the concentration of 20 different neural metabolites, including GABA and glutamate, from a 2×2×2cm^3^ voxel placed just anterior of the anodal electrode (Fig. 3D). Consistent with previous literature, application of anodal tDCS was accompanied by a significant reduction in the concentration of GABA relative to baseline (‘baseline’ > ‘tDCS’, t_19_= 1.97, p=0.016, Fig. 3E). As a consequence, block 2 of the scan task was performed in a state of EI imbalance, where excitation outweighed inhibition. The reduction in GABA was not sustained to the period after the task (‘baseline’ > ‘post-task’, p>0.05, t_19_=0.22, p=0.414, Fig. 3E). In addition to this change in GABA we also observed a significant increase in the concentration of glutamate, but only after the second task session (‘baseline’ < ‘post-tDCS’, t_19_=2.26, p=0.018; Fig. 3F). This change in glutamate may be attributed to participants performing block 2 of the scan task, and doing so in a state of EI imbalance (see Supplementary Table 1 for list of all measured metabolites).

### Measuring associative memories using cross-stimulus suppression

To test whether neocortical inhibition protects memories from interference, we assessed evidence for neural memory interference during the period of EI imbalance. To measure memory interference in neocortex we first sought to index neocortical co-activation between representations for different memory elements. We took advantage of fMRI repetition suppression which relies on the fact that neurons show a relative suppression in their activity in response to repeated presentation of a stimulus to which they are sensitive (Miller et al., 1991; Sawamura et al., 2005). While typically used to access sub-voxel representations for single stimuli (Grill-Spector et al., 2006), ‘cross-stimulus’ suppression can be used to index the relative co-activation or overlap between representations coding for two different stimuli (Barron et al., 2016b). Here, we hypothesized that we could use cross-stimulus suppression to measure the relative co-activation between pairs of elements from memory 1 and memory 2, using the background colour to provide a contextual cue. We contrasted the BOLD response for each pair of stimuli where suppression was expected against the BOLD response to a control pair where suppression was not expected, thus controlling for attentional effects (Fig. 3G). The ring topology of memory 1 and 2 provided an efficient way to ensure that each stimulus contributed to both trials where suppression was expected (directly associated stimuli in one or both contexts) and control trials where suppression was not expected (stimuli separated by up to three associations in both contexts).

### Otherwise dormant associative memories are unmasked during periods of EI imbalance

Having induced a transient period of EI imbalance, we used cross-stimulus suppression to assess the effect of reduced GABAergic tone on neocortical associative memories. Previously, we have shown that manipulating GABAergic tone in this way increases cross-stimulus suppression between directly associated stimuli in aLOC (Fig. 3C) (Barron et al., 2016a). Here we sought to first replicate this result, before going on to assess the effect of reduced GABAergic tone on memory interference.

Given that we were aiming to replicate our previous findings (Barron et al., 2016a), we extracted parameter estimates from an independently defined region of interest (ROI) in aLOC, taken from our previous data set (Fig. 3H). Within this ROI we observed a significant increase in cross-stimulus suppression during anodal tDCS (Block 2 − Block 1), across memory 1 and 2 (t_23_=1.73, p=0.049, Fig. 3I), and for memory 2 (t_23_=2.31, p=0.016, Fig. 3J-K), but not memory 1 alone (p>0.05, Supplementary Fig. 4A). Furthermore, during brain stimulation (Block 2) we observed significant cross-stimulus suppression within this ROI for memory 1 and memory 2 together (t_23_=2.31, p=0.015, Fig. 3I), and for memory 2 (t_23_=2.91, p=0.004, Fig. 3J-L), but not memory 1 alone (p>0.05, Supplementary Fig. 4A). Significant cross-stimulus suppression for directly associated elements of memory 2 could also be observed in Block 2 within a 10mm sphere centered on the peak of the average anodal tDCS electrode location (Fig. 3M, see Methods), correcting for multiple comparisons using a small volume correction (SVC) method (t_23_=4.17, p=0.010, peak-level family-wise error (FWE) corrected). Thus, replicating our previous findings (Barron et al., 2016a), these results show that reducing GABAergic tone increases the expression of otherwise dormant associative memories, particularly those formed more recently.

**Figure 4:**
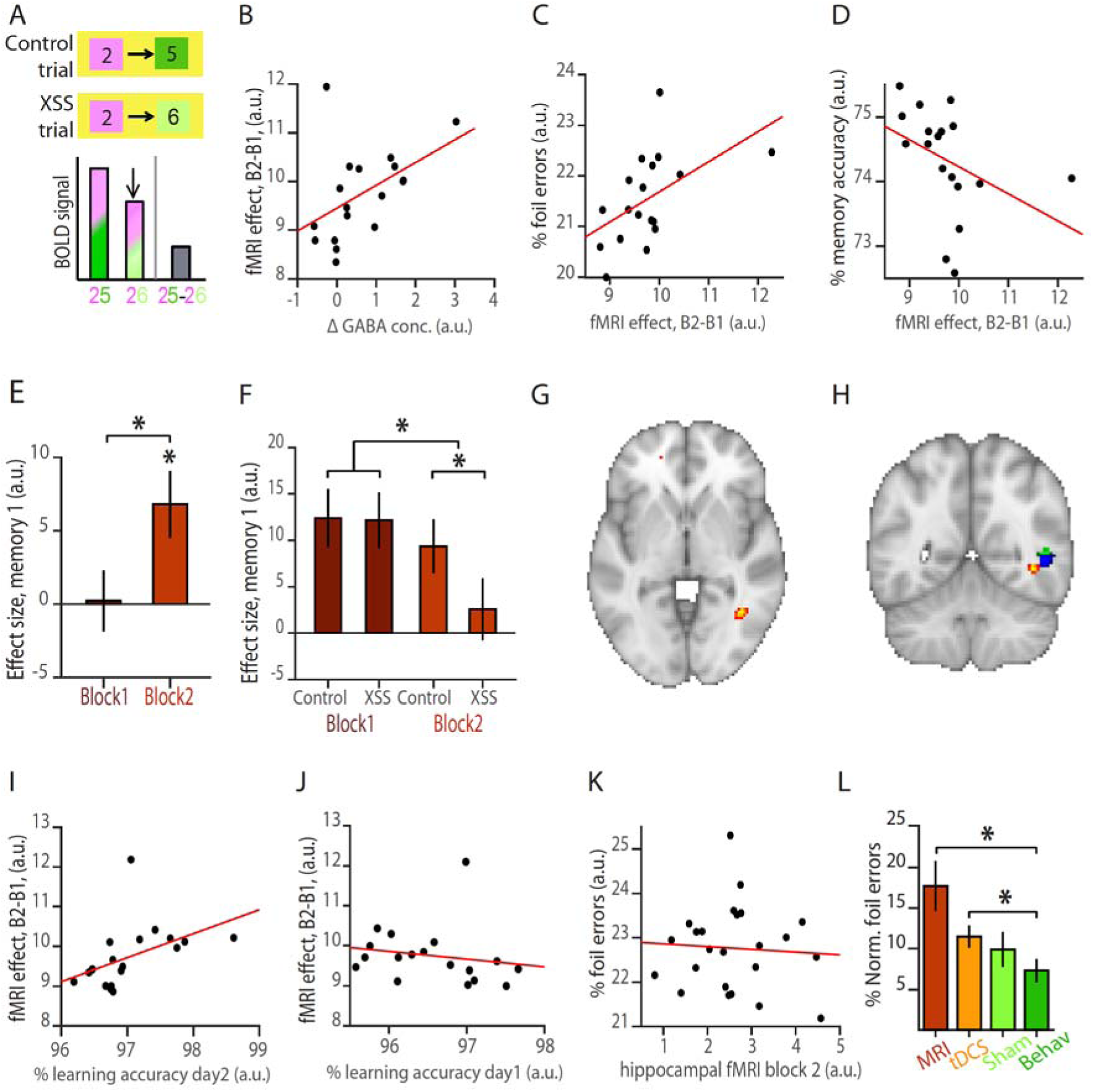
Memory interference for more remote memories increases with brain stimulation. **A**) When participants performed the scan task in EI imbalance, we predicted an increase in cross-stimulus suppression (‘XSS’) on trials where participants observed pairs of stimuli that were unassociated in the current context but directly associated in the alternative context, relative to control trials where participants observed pairs of stimuli that were unassociated in both contexts. This difference between ‘Control’ and ‘XSS’ trials could be measured using the BOLD signal and used as an index for neural memory interference. **B)** Across participants, the change in GABA concentration (‘Before tDCS’ - ‘During tDCS’) positively predicted the increase in fMRI cross-stimulus suppression used to measure interference in memory 1 and 2 (‘Control’ - ‘XSS’ for Block 2 - Block 1) (B indicates ‘Block’, Spearman correlation: r_17_=0.55, p=0.021, after accounting for changes in glutamate, see Methods). **C)** Across participants, the cross-stimulus suppression index for memory interference (‘Control’ - ‘XSS’ for Block 2 - Block 1) positively predicted the percentage of foil errors participants made on the surprise memory test on day 3 (Spearman correlation: r_17_=0.58, p=0.013). **D)** Across participants, the cross-stimulus suppression index for memory interference (‘Control’ - ‘XSS’ for Block 2 - Block 1) negatively predicted average memory accuracy on the surprise memory test on day 3 (Spearman correlation: r_17_=-0.67, p=0.003). **E)** Within an ROI defined from the peak average tDCS electrode location shown in Fig. 3D, extracted parameter estimates for memory 1 (shown: mean ±SEM) revealed a significant increase in the fMRI cross-stimulus suppression measure for memory interference (‘Control’ - ‘XSS’, as shown in ***A)*** from Block 1 to 2 and during Block 2 alone (‘Control’ - ‘XSS’ for Block 2 - Block 1: t_23_=2.75, p=0.006; ‘Control’ - ‘XSS’ for Block 2: t_23_=2.70, p=0.006). **F)** Extracted parameter estimates from *E* split into the ‘Control’ and ‘XSS’ conditions, as described in *A* (shown: mean ±SEM). **G)** T-statistic map for cross-stimulus suppression index for neural memory interference during Block 2 between unassociated stimuli in memory 1 that are directly associated in memory 2, relative to pairs of stimuli that are unassociated in both memories, thresholded at p<0.01 uncorrected for visualization. **H)** Illustrating the anatomical proximity between the effects shown in Fig. 4G (red), Fig. 3L (blue) and previously acquired data set shown in Fig. 3C (green). **I)** Across participants, learning accuracy on memory 2 (day 2) positively predicted the cross-stimulus suppression index for neural memory interference (Spearman correlation: r_17_=0.68, p=0.003). **J)** Across participants, learning accuracy on memory 1 (day 1) showed a negative trend with the cross-stimulus suppression index for neural memory interference (Spearman correlation: r_17_=-0.47, p=0.053). **K)** During application of anodal tDCS in the second scan task (Block *2*), the hippocampal BOLD response to trials where there was opportunity for memory interference did not predict behavioural measures of memory interference (Pearson correlation: r_2_3=-0.06, p=0.764). **L)** Both groups of participants (see Methods) who received tDCS (‘MRI’ and ‘tDCS’) showed higher normalised foil errors on the surprise memory test on day 3, relative to participants who received no intervention (‘Behav’) (two-sample t-test: t_44_=2.89, p=0.006). However, there was no difference in the percentage of foil errors made by participants who received ‘tDCS’ and ‘Sham’ (two-sample t-test: t_38_=0.66, p=0.515). Normalised foil errors were defined as the percentage of foil errors on foil trials subtracted by the percentage of non-foil errors on foil trials.

To further assess the relationship between the down-regulation of GABA and the change in cross-stimulus suppression we measured the correlation between these two variables. The drop in GABA during application of anodal tDCS positively predicted the increase in cross-stimulus suppression between directly associated stimuli in memory 1 and 2 (Fig. 3N, r_17_=0.52, p=0.028, after accounting for changes in glutamate, see Methods). Notably, the variation in the drop in GABA observed across participants provided a stringent framework in which to assess the effect of EI imbalance on cross-stimulus suppression. Participants with a small drop in GABA provided effective parametric control for participants with a larger drop in GABA, thus mitigating the need for a separate sham condition.

These results replicate our previous findings, showing that associative memories are re-expressed following down-regulation of neocortical GABA. This implies that associative memories are stored in neocortex in EI balanced ensembles, where, if left unperturbed, neocortical inhibition acts to quench memory expression.

### Memory interference increases during periods of EI imbalance

Having replicated our previous findings (Barron et al., 2016a), we went on to investigate whether neocortical inhibition plays a critical role in protecting against memory interference. Capitalizing on the inter-subject variability to the anodal tDCS manipulation (Fig. 3E), we predicted a two-way relationship between the drop in GABA, neural measures of memory interference and behavioural measures of memory interference: if neocortical inhibition protects memories from interference, the drop in GABA should predict neural measures of memory interference, which should in turn predict behavioural measures of memory interference.

As neural memory interference manifests as activation of a relational neighbour from the alternative, inappropriate memory, we sought to index this unwanted activation using cross-stimulus suppression. To this end, we identified trials during the scan task where participants were shown two stimuli that were unassociated given the memory indicated by the contextual cue, but directly associated in the alternative memory. During periods of EI imbalance we predicted an increase in cross-stimulus suppression on these trials, relative to trials where the presented stimuli were indirectly associated in both memory 1 and 2 (Fig. 4A). Thus, an *increase* in this cross-stimulus suppression measure provided a proxy for an *increase* in neural memory interference. Using this measure of memory interference we assessed evidence for the predicted two-way relationship between the drop in concentration of GABA, neural memory interference and behavioural memory interference.

Firstly, we considered the relationship between the drop in GABA during application of anodal tDCS and the increase in neural memory interference from the first to the second fMRI scan task. Across participants, the drop in GABA positively predicted the increase in neural memory interference measured using cross-stimulus suppression across memory 1 and 2 (r_17_=0.55, p=0.021, Fig. 4B, after accounting for changes in glutamate, see Methods). Cross-stimulus suppression measured from participants with minimal change in the concentration of GABA thus parametrically controlled for participants where a larger change was observed. This result suggests that interference between overlapping memories is predicted by EI imbalance.

Secondly, we considered the relationship between neural and behavioural memory interference. Taking behavioural performance from the surprise memory test performed after the scan, we predicted a positive relationship between neural memory interference and the percentage of foil errors on the memory test, but a negative relationship between neural memory interference and overall accuracy on the memory test. Consistent with these predictions, our cross-stimulus suppression index for neural memory interference (Block 2 – Block 1) positively predicted the percentage of foil errors, and negatively predicted overall behavioural memory accuracy (fMRI vs. foil errors: r_17_=0.58, p=0.013, Fig. 4C; fMRI vs. overall accuracy: r_17_=-0.67, p=0.003, Fig. 4D; after accounting for differences in learning and changes in GABA and glutamate, see Methods). In summary, participants who showed greater cross-stimulus suppression during periods of EI imbalance also made more errors.

Together these results suggest that a reduction in neocortical GABAergic tone leads to an increase in neural memory interference which manifests in behaviour as an increase in memory errors. While this two-way relationship capitalizes on the variability observed across participants, we next asked whether there was a main effect of anodal tDCS on neural memory interference. Using cross-stimulus suppression as a proxy for memory interference, we predicted an overall increase in neocortical memory interference during the application of anodal tDCS. Furthermore, given that a reduction in neocortical GABA resulted in pronounced re-expression of more recent associations in memory 2 (Fig. 3J-L), we predicted memory interference would manifest in memory 1 due to expression of associations in memory 2 intruding or overriding the appropriate expression of associations in memory 1.

To maximize sensitivity to the effect of anodal tDCS, we tested for memory interference using cross-stimulus suppression within an ROI defined from the peak anodal tDCS electrode location, averaged across all participants (Fig. 3D, see Methods). Within this ROI, across both memory 1 and 2 we observed a trend towards an increase in cross-stimulus suppression during application of anodal tDCS (Block 2 > Block 1, t_23_=1.40, p=0.089, Supplementary Fig. 4B). However, consistent with our prediction, for memory 1 but not memory 2 there was a pronounced increase in our cross-stimulus suppression measure of memory interference (Block 2 > Block 1, memory 1: t_23_=2.75, p=0.006 Fig. 4E-F; memory 2: p>0.05, Supplementary Fig. 4C). To confirm that memory interference was observed during application of anodal tDCS, we also assessed effects in block 2 alone. Within the same ROI we again observed significant cross-stimulus suppression for memory 1 but not memory 2 (memory 1: t_23_=2.70, p=0.006, Fig. 4E-G), and within a 10mm sphere centered on the peak of the average anodal tDCS electrode location (memory 1: t_23_=3.60, p=0.027, peak-level family-wise error (FWE) corrected using a small volume correction (SVC) method with ROI shown in Fig. 3M). Critically, this cross-stimulus measure of interference in memory 1 was anatomically proximal to the re-expression of directly associated memories in memory 2 reported above (Fig. 4H).

These results suggest that re-expression of directly associated stimuli in memory 2 leads to interference with overlapping, but contextually distinct, associations in memory 1. In a final analysis, we asked whether the differential strength of memory 1 and 2 at encoding also predicts neural memory interference during periods of EI imbalance. We found that participant’s average learning accuracy for associations in memory 2 positively predicted the cross-stimulus suppression measure for memory interference (r_17_=0.68, p=0.003, Fig. 4I, after accounting for differences in learning on day1, memory accuracy and changes in GABA and glutamate, see Methods), while a trend towards the reverse relationship was observed for memory 1 (r_17_=-0.47, p=0.053, Fig. 4J, after accounting for differences in learning, memory accuracy and changes in GABA and glutamate, see Methods). Therefore, participants that weakly encoded memory 1 but strongly encoded memory 2 were more prone to memory interference. Together with results above, this suggests that interference between two memories during periods of EI imbalance can be predicted by the relative strength of the memories at encoding.

### The interplay between the hippocampus and neocortex

These data suggest that in addition to context-dependent relational codes in the hippocampus, neocortical inhibition also appears to play a key role in protecting memories from interference. To assess the interplay between the hippocampal and neocortical mechanisms, we reconsidered the relationship between our neural and behavioural measures of memory interference. We noted that behavioural performance on the surprise memory test after the scan session was predicted by both the hippocampal BOLD signal prior to anodal tDCS (Fig. 2H), and the change in neocortical cross-stimulus suppression observed during anodal tDCS (Fig. 4C-D). We asked whether hippocampal BOLD *during* anodal tDCS (block 2) also predicted behavioural performance. Unlike hippocampal BOLD prior to anodal tDCS (block 1), we observed no relationship between hippocampal BOLD during tDCS (block 2) and behavioural performance (hippocampal BOLD block 2 vs. foil errors: r_23_=-0.06, p=0.764, Fig. 4K, after accounting for differences in learning). Furthermore, this correlation between block 2 hippocampal BOLD and behaviour was significantly different from the correlation observed between block 1 hippocampal BOLD and behaviour (difference in correlation coefficient, block 1 vs block 2, permutation test: p=0.032, Supplementary Fig. 4D, see Methods). This suggests that in the absence of brain stimulation, the degree to which irrelevant associative links are activated in the hippocampus predicts memory interference. But, when neocortical GABAergic tone is reduced, the degree to which irrelevant associative links are activated in the neocortex but not hippocampus predicts memory interference. The hippocampus and neocortex thus appear to employ distinct mechanisms to mediate memory interference.

### Inducing behavioural memory interference using brain stimulation

In a final set of experiments we asked whether application of anodal tDCS alone might be sufficient to induce behavioural measures of memory interference. To this end we repeated the experiment in three additional groups of participants: (2) anodal tDCS or (3) sham-anodal tDCS (delivered using a double-blind set-up, see Methods), or (4) no intervention. These three additional groups of participants performed the same set of tasks as participants receiving MRI (group 1), but outside the scanner. On the day 3 surprise memory test we observed a significant difference between groups in mean accuracy and the percentage of normalised foil errors using a one-way ANOVA (mean accuracy: F_85_=6.54, p<0.001, Supplementary Fig. 2C; normalised foil errors: F_85_=4.39, p =0.007, Fig. 4L). Post-hoc t-tests revealed a significant difference between participants who received both anodal tDCS and MRI compared to participants who did not receive any intervention (group 1, ‘MRI’, vs. group 4, ‘Behav’, foil errors: t_44_=2.89, p=0.006, Fig. 4L), and for participants who received anodal tDCS compared to participants who did not receive any intervention (group 2, ‘tDCS’, vs. group 4, ‘Behav’, foil errors: t_38_=2.30, p=0.027, Fig. 4L). For participants who received sham-stimulation, there was no significant difference in behavioural performance compared to participants who did not receive any intervention (group 3, ‘Sham’, vs. group 4, ‘Behav’, foil errors: t_38_=0.18, p=0.860, Fig. 4L). While these results suggest that anodal tDCS increased memory interference at the behavioural level, there was notably no significant difference in behavioural performance between participants who received anodal tDCS and those who received sham stimulation (group 2, ‘tDCS’, vs. group 3, ‘Sham’,: foil errors: t_38_=0.66, p=0.515, Fig. 4L), suggesting that while anodal tDCS can induce behavioural memory interference, the expectation of anodal tDCS has a similar effect on some participants. Therefore, rather than mere application of brain stimulation, the change in the concentration of GABA and neural measures of memory interference are necessary to reliably predict behavioural measures of memory interference.

## Discussion

Our past experiences often overlap in their content, but can nevertheless be selectively recalled without interference from other memories. Here, we investigated the neural mechanisms that help protect memories from interference. By training human participants to encode two context-dependent overlapping memories, memory 1 and 2, we investigated the neural mechanisms that mitigate against memory interference. We reveal evidence for two distinct mechanisms. The first mechanism involves the hippocampus, where memories are organised according to context-dependent relational codes. The second mechanism involves neocortical inhibition, which protects against unwanted co-activation between neocortical representations. We discuss these two mechanisms in turn, before considering how they may interact.

In the hippocampus we observed an increase in hippocampal BOLD signal when participants observed pairs of stimuli that had different relative positions across memory 1 and 2. In the absence of brain stimulation this BOLD signal predicted participants’ performance on a surprise memory test completed after the scan. When we investigated the nature of the underlying hippocampal representations, we found that stimuli with different relational positions across memory 1 and 2 had more distinct representations across memory 1 and 2. These findings are in agreement with evidence that the hippocampus represents contextual information (Butterly et al., 2012; McKenzie et al., 2014). However, they also suggest that contextual representations are organised using a relational code. Critically, this reconciles two views of the hippocampus: one in which contextual representations help orthogonalize competing memories (McKenzie et al., 2014), and a second where overlapping memories are integrated into relational structures or schemas (Dabaghian et al., 2014; Eichenbaum, 2017; Garvert et al., 2017; Konkel and Cohen, 2009). Therefore, for two overlapping but context-dependent memories, divergent representations may be observed for items or stimuli that have different relational positions, while similar representations may be observed for items or stimuli that have similar relational positions. Interestingly, this account is consistent with the idea that the hippocampus represents a successor representation where stimuli that predict different future states have more distinct representations (Garvert et al., 2017; Momennejad et al., 2017; Stachenfeld et al., 2017).

While the hippocampus may help minimize interference between overlapping memories using context-dependent relational representations, the sensory neocortex appears to employ a different mechanism. By down-regulating the concentration of neocortical GABA using anodal tDCS (Barron et al., 2016a; Kim et al., 2014; Stagg et al., 2009), here we show that during periods of EI imbalance, neocortical memory interference increases. To quantify neural memory interference we used ultra-high field 7T MRI to measure cross-stimulus suppression, a proxy for representational similarity between different elements of the memories (Barron et al., 2016b; Krekelberg et al., 2006). We show that the drop in GABA quantified using MRS predicts our neural measure of memory interference, which in turn predicts behavioural measures of memory interference. This two-way relationship reveals a key role for neocortical inhibitory engrams in protecting against interference.

While these findings suggest that inhibitory engrams are critical for stable memory storage, they also raise a number of questions regarding the formation of inhibitory engrams and the accompanying timescale of this process. In the rodent primary auditory cortex, changes in the strength of excitatory connections are accompanied by inhibitory rebalancing after approximately 90 minutes (Froemke et al., 2007). This implies that a ‘critical period’ of EI imbalance and memory instability occurs between initial learning and the formation of inhibitory engrams. Consistent with this hypothesis, a transient period of memory instability has been reported immediately after learning, during which memories may be integrated with existing knowledge that share abstract or higher-level properties (Mosha and Robertson, 2016). This integration is facilitated by offline reactivation and coordinated interactions between hippocampal and neocortical engrams (Mosha and Robertson, 2016; Preston and Eichenbaum, 2013; Schlichting and Preston, 2014).

While this opportunity to integrate newly encoded memories with existing knowledge has clear advantages, the relative instability of memories during this ‘critical period’ makes them vulnerable to interference. This trade-off between integration and interference may determine the transient nature of the ‘critical period’. Indeed, if sufficient time is left between acquisition of memories that are overlapping or share a common structure, integration is no longer observed (Mosha and Robertson, 2016), nor is interference, as shown here. In addition to time, other factors such as overlearning also appear to terminate the ‘critical period’ (Mosha and Robertson, 2016) by restoring EI balance with a shift from glutamate-dominated excitation to GABA-dominated inhibition (Shibata et al., 2017).

By combining ultra-high field 7T fMRI with MRS, brain stimulation and behavioural measures, the protocol described here illustrates how macroscopic measures of the human brain can be used to index micro-circuit processes. This has notable translational value for clinical populations where microcircuit disruption is not readily amendable to investigation, particularly conditions that have been attributed to disturbances in EI balance. For example, in schizophrenia delusions and hallucinations have been attributed to perturbed inhibitory gating (Vogels and Abbott, 2007; Yizhar et al., 2011), while memory loss and confusion in early stage dementia have been associated with hyperactivity (Busche and Konnerth, 2016). The data presented here may be considered a model for these clinical phenotypes, where neocortical EI imbalance causes unwanted reactivation of irrelevant memories that may have overlapping excitatory engrams with those activated by incoming stimuli. In the absence of appropriate inhibitory regulation, otherwise latent memory engrams are activated in an uncontrolled manner causing confusion or hallucinations. The protocol implemented here thus provides a basis from which to further explore mechanisms responsible for memory impairment in clinical conditions that report evidence for EI imbalance.

Finally, we consider the interplay between the hippocampal and neocortical mechanisms for mitigating against memory interference. When we reduced the concentration of neocortical GABA using anodal tDCS, interestingly the hippocampal BOLD signal no longer predicted behavioural performance on the surprise memory test. This suggests that inhibitory gating in the neocortex may influence context-dependent hippocampal representations. Although beyond the scope of this investigation it is also interesting to speculate about the reverse relationship, the influence of context-dependent relational hippocampal codes on neocortical memory engrams. One possibility is that analogous to cholinergic modulation of neocortical interneurons observed during context-dependent behaviour (Kuchibhotla et al., 2017), the hippocampus may mediate selective release of neocortical memory engrams by targeting neocortical inhibition (Barron et al., 2017). This interaction between the hippocampus and neocortex may facilitate selective reactivation of neocortical representations during memory recall, providing an index for distributed memory engrams (Teyler and Rudy, 2007).

## Supplementary Figures

**Supplementary Figure 1.**
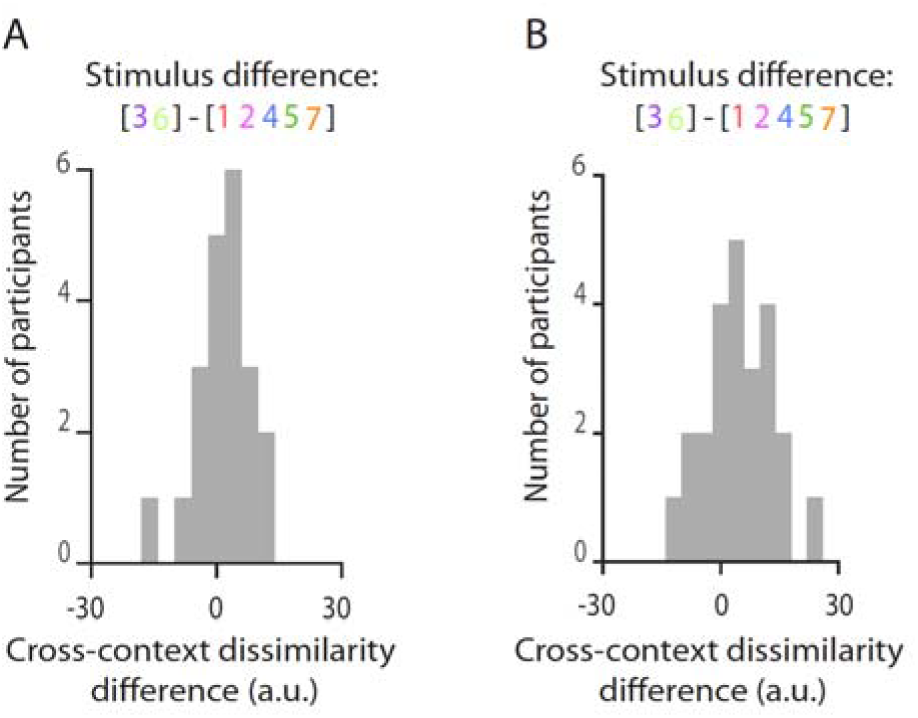
**A**) Histogram showing the difference in average hippocampal dissimilarity between each stimulus in one memory and all stimuli in the other memory, across all participants. On average, the dissimilarity difference was greater for stimuli that had different relative positions across memory 1 and 2 (stimuli 3 and 6) compared to stimuli that had the same relative position across memory 1 and 2 (stimuli 1, 2, 4, 5, 7) (Wilcoxon sign rank test: Z_23_=1.97, p=0.048). **B)** Histogram showing the difference in average hippocampal dissimilarity between each stimulus in memory 2 and all stimuli in memory 1 across all participants. On average, the dissimilarity difference was greater for stimuli that had different relative positions across memory 1 and 2 (stimuli 3 and 6) compared to stimuli that had the same relative position across memory 1 and 2 (stimuli 1, 2, 4, 5, 7) (Wilcoxon sign rank test: Z_23_=2.51, p= 0.012).

**Supplementary Figure 2.**
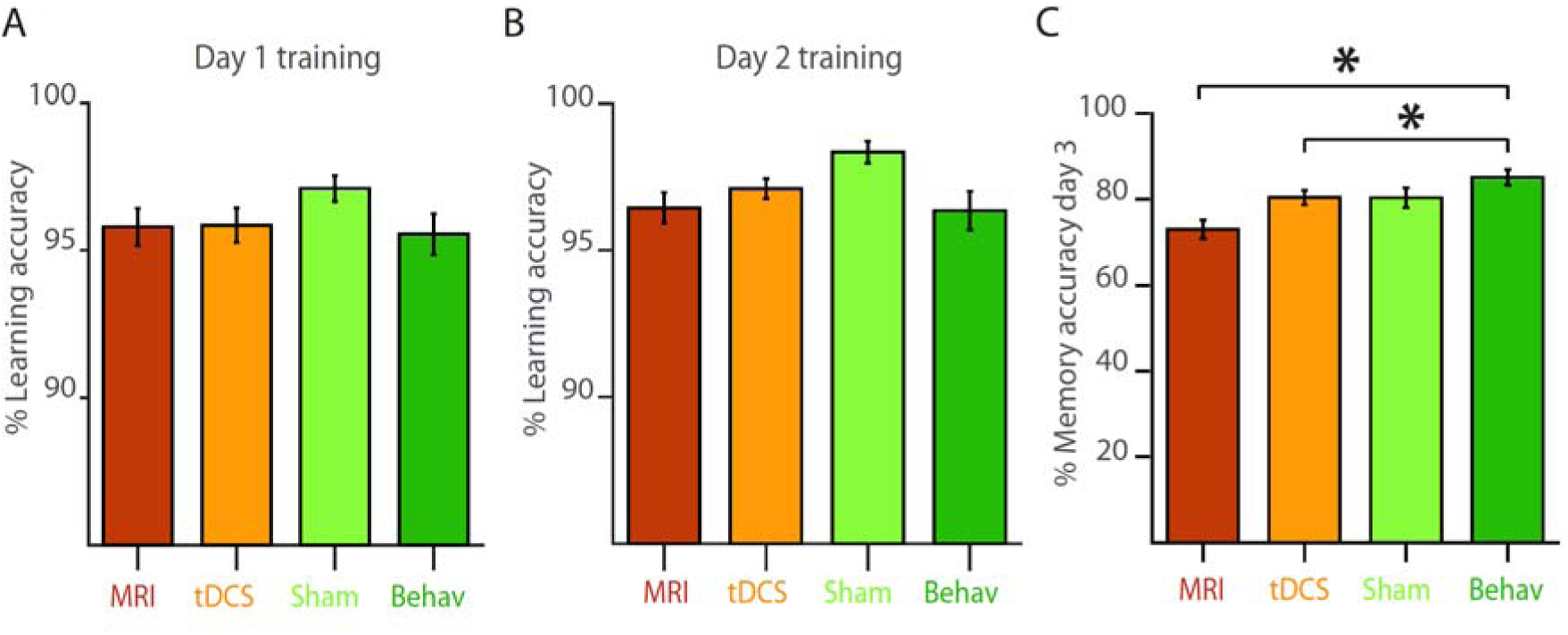
**A**) Percentage learning accuracy on the training task used on day 1 (Fig. 1D) for the four different groups of participants (see Methods), averaged across trials on each participant’s highest performing task block. **B)** Percentage learning accuracy on the training task used on day 2 (Fig. 1E) for the four different groups of participants (see Methods), averaged across trails on each participant’s highest performing task block. **C)** Accuracy on the surprise memory test on day 3 for all four groups of participants. Participants who received tDCS (‘MRI’ and ‘tDCS’) showed lower memory accuracy relative to participants who received no intervention (‘Behav’) (t_44_=4.11, p<0.001). However, there was no difference in performance between the ‘tDCS’ and ‘Sham’ groups (t_38_=0.02, p=0.986).

**Supplementary Figure 3.**
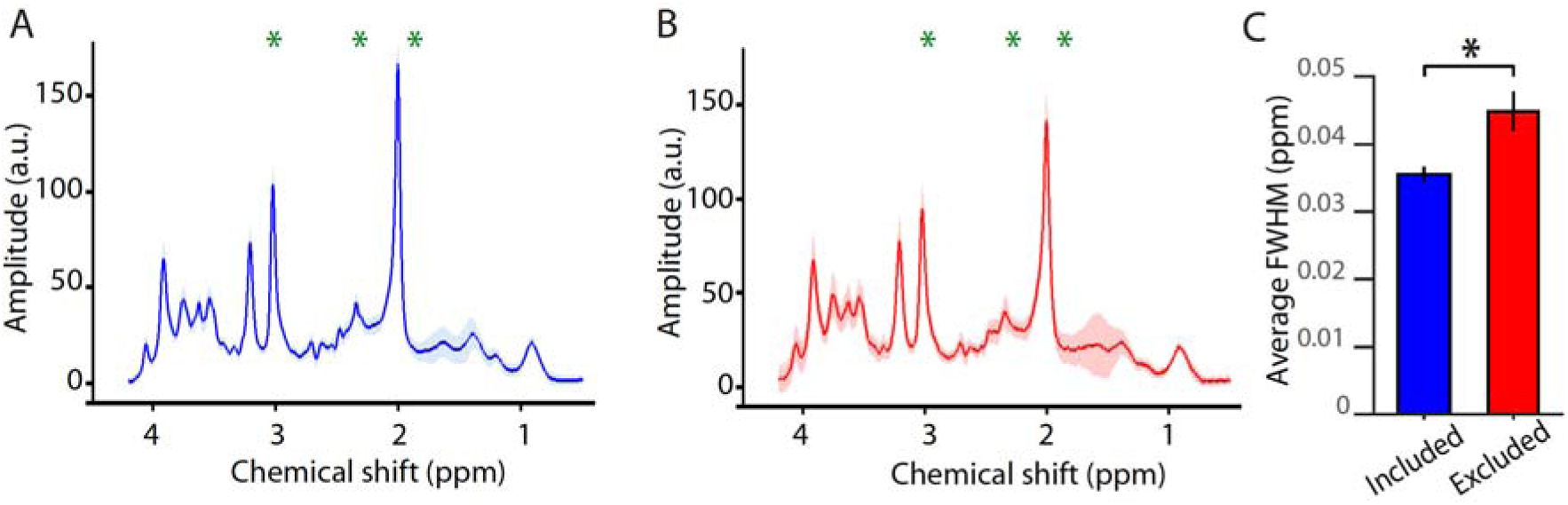
**A**) Average spectra for all participants data included in MRS analysis. Chemical shifts of the three GABA peaks are indicated using green stars. **B)** Average spectra for those participants who were rejected from the MRS analysis. Data from these participants were noisy, and had lipid contamination in the region 1.9-0.5ppm (i.e. in the region of the lowest GABA peak). This resulted in either inestimable or highly unreliable GABA estimates. Chemical shifts of the three GABA peaks are indicated using green stars. **C)** Relative to participants included in the MRS analysis (shown in ***A)***, those participants rejected from the MRS analysis (shown in ***B*)** had broader linewidth, estimated using full-width at half maximum (FWHM) using LCModel (two-sample t-test: t_24_=3.33, p=0.001).

**Supplementary Figure 4.**
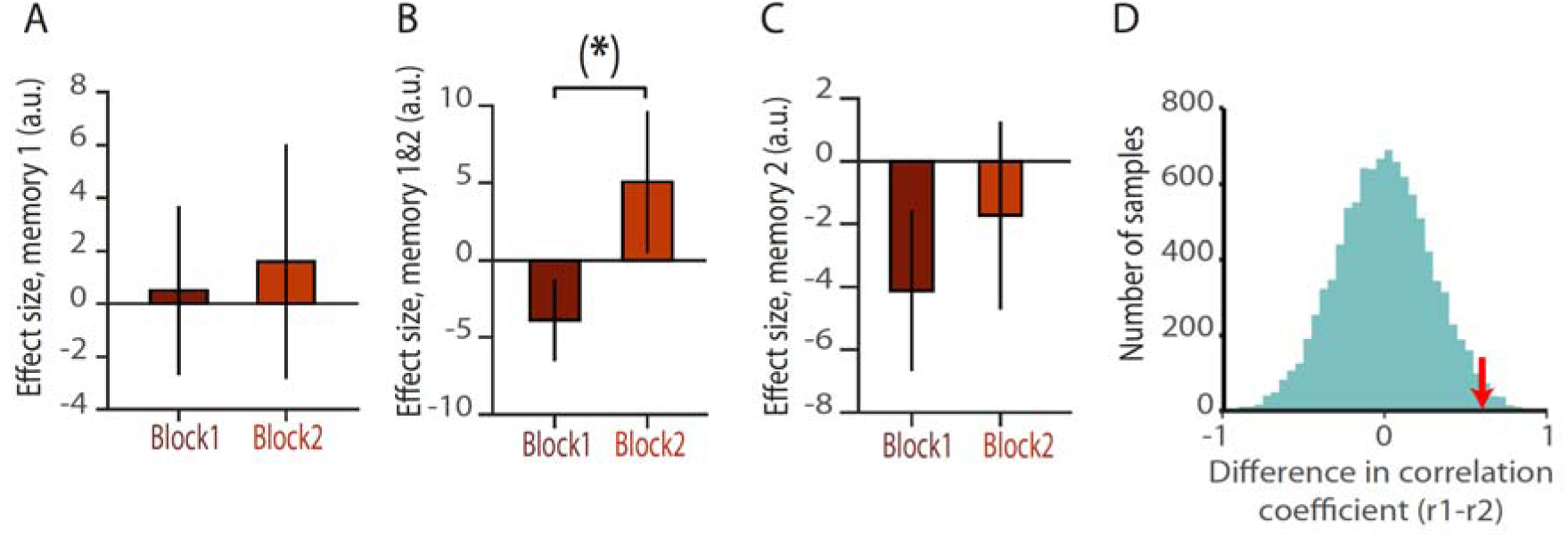
**A**) Cross-stimulus suppression was used to index the change in expression of directly associated stimuli in memory 1 before and during application of tDCS. Unlike for memory 2 (Fig. 3J-L), no significant change was observed for memory 1 (p>0.05, shown: mean ±SEM). **B)** Cross-stimulus suppression was used to index the change in memory interference before and during application of tDCS. Across memory 1 and 2 there was a trend towards an increase in the cross-stimulus suppression (paired t-test: t_24_=1.40, p=0.087). **C)** Cross-stimulus suppression was used to index the change in memory interference in memory 2 before and during application of tDCS. Unlike for memory 1 (Fig. 4E-F), no significant change was observed for memory 2 (p>0.05). **D)** During the first but not the second scan task hippocampal BOLD predicted subsequent behavioural performance on the surprise memory test (Fig. 2H and Fig. 4K). To assess the significance of the difference in correlation coefficient, a null distribution of 10,000 samples was estimated using a permutation test (see Methods). Here, the null distribution can be observed in green and the difference in correlation indicated by the red arrow.

## Supplementary Tables

**Supplementary Table S1.** (related to Fig. 3-4) | Average concentration of all metabolites measured using MRS. For each metabolite, the concentration was measured relative to creatine and then averaged across participants. As reported in the main text, a significant decrease in the concentration of GABA was observed during tDCS, and a significant increase in the concentration of glutamate observed after the second task block (Fig. 3E-F). Of the other metabolites measured (n=17), only one showed a significant change in concentration across the three MRS measurements: the concentration of aspartate significantly decreased after the second task block (‘before tDCS’ - ‘post-task’, t_19_=4.29, p<0.001).

| Metabolite | Before tDCS | During tDCS | Post-task |
| --- | --- | --- | --- |
| Alanine | 1.39 | 1.52 | 1.31 |
| Ascorbate | 1.11 | 1.29 | 1.22 |
| Aspartate | 3.18 | 2.78 | 2.26 |
| Glycerophosphorylcholine | 0.93 | 0.86 | 0.94 |
| Phosphorylcholine | 0.75 | 0.83 | 0.76 |
| Creatine | 4.96 | 4.97 | 4.91 |
| Phosphocreatine | 3.04 | 3.03 | 3.09 |
| GABA | 4.77 | 4.18 | 4.53 |
| Glucose | 2.22 | 2.31 | 2.31 |
| Glutamine | 7.30 | 7.36 | 7.04 |
| Glutamate | 9.26 | 9.39 | 9.75 |
| Glutathione | 0.89 | 0.77 | 0.79 |
| Inositol | 7.27 | 7.19 | 7.19 |
| Lactate | 1.10 | 1.18 | 0.89 |
| N-acetylaspartate (NAA) | 13.25 | 13.19 | 13.28 |
| N-acetylaspartylglutamate (NAAG) | 2.34 | 2.29 | 2.31 |
| Phosphoethanolamine | 1.91 | 1.97 | 1.95 |
| Scyllo-Inositol | 0.18 | 0.17 | 0.17 |
| Taurine | 0.65 | 0.53 | 0.51 |

## Acknowledgements

R.S.K. is supported by an EPSRC/MRC-funded studentship. T.E.B. is supported by a Wellcome Trust Senior Research Fellowship (WT104765MA), together with a James S. McDonnell Foundation Award (JSMF220020372). H.C.B. is supported by a Junior Research Fellowship from Merton College (University of Oxford) and the John Fell Oxford University Press Research Fund (Grant 153/046). The Wellcome Centre for Integrative Neuroimaging is supported by core funding from the Wellcome Trust (203139/Z/16/Z).

## Author contribution

All of the authors contributed to the design of the study and preparation of the manuscript. R.K., U.E.E., A.C.P. and H.C.B. acquired the data and R.K., U.E.E. and H.C.B. analysed the data.

## Methods

### Participants

88 healthy volunteers participated in the study (group 1: “MRI” with tDCS, n = 27, mean age of 24.1, 14 females; group 2: “tDCS”, n = 20, mean age of 21.9, 18 females; group 3: “Sham”, n = 20, mean age of 24.1, 9 females; group 4: “Behav”, n =21, mean age of 23.2, 11 females). All experiments were approved by the Oxford University ethics committee (reference number ref R43594/RE001). All participants gave informed written consent.

In group 1, one participant was excluded due to a fault on the scanner which prevented data acquisition in the second half of the scan session. In group 4, one participant was excluded after revealing that they had an arteriovenous malformation in the cerebellum.

### Behavioural training

All behavioural tasks were coded in Matlab 2016b using Psychtoolbox (version 3.0.13). Seven different stimuli were presented to the participant, referred to as 1:7. Stimuli were rotationally invariant gray shapes (Fig. 1A), which were observed in one of four possible rotations, with each rotation separated by 90° (as described in (Barron et al., 2016a)). The experiment was conducted across three days. On the first day participants learned seven bidirectional associations between the seven stimuli. The set of associations could be arranged in a ring structure (Fig. 1B), where each stimulus was associated with two other stimuli (1 with 2, 2 with 3, etc. and 7 with 1). Participants were not explicitly made aware of the ring structure and instead learned the associations using a training task. Stimulus allocation within the ring structure was randomised across participants using Matlab’s random number generator.

The training task first involved passive exposure to the seven associations with each pair presented for 2 s duration. This passive phase of the task was then followed by a three-alternative forced-choice task (Fig. 1D-E). On each trial of the three-alternative forced-choice task, one of the seven stimuli was shown as a probe stimulus for 1 s before three choice stimuli were presented in randomised positions across the screen. Stimuli were presented against a background colour (blue or yellow), used to provide a contextual cue. The three choice stimuli included one stimulus to which the probe stimulus was associated, and two stimuli to which the probe stimulus was not associated. Participants were instructed to select the correctly paired stimulus as fast as possible, without compromising their accuracy, using the appropriate keyboard button ‘b’, ‘n, or‘m’. If participants failed to make a response within 3 s they received an on-screen message indicating that they were too slow. Participants received feedback for each choice, where the probe stimulus together with the correctly paired choice stimulus were presented for 1.5 seconds. For each correct response, participants were assigned 50p. At the end of each task block (100 trials) three percent of trials were randomly selected and participant received the sum total reward from these trials. To qualify for the second day of the experiment, participants were required to obtain at least 90% accuracy on all of the seven associations and to complete at least five blocks of the tasks.

On the second day, participants again learned seven associations between the seven stimuli, however the position of the stimuli within the implicit ring structure was altered relative to the first day. In particular, stimuli ‘3’ and ‘6’ were switched, resulting in four new associations and three associations that remained the same across days (Fig. 1C). To indicate this change in the implicit associative structure, the background colour of the screen (blue or yellow) was changed from day 1 to day 2. The colour assigned to day 1 and 2 was randomised across participants. To learn the new arrangement of stimuli, participants underwent the same training task as that used on day 1, including both the passive task and the three-alternative forced-choice tasks (Fig. 1E). To qualify for the third day of the experiment, participants were required to obtain at least 90% accuracy on all of the seven associations in the day 2 ring and to complete at least five blocks of the task.

On the third day of the experiment, participants were required to perform the fMRI scan task (see below, Fig. 1F). The fMRI scan task was performed inside the scanner for group 1, but outside the scanner for groups 2-4. Immediately after exiting the scanner (group 1), or immediately after the scan task (groups 2-4), participants were given a surprise memory test designed to assess participants’ memory for the associations learned on both day 1 and day 2. The memory test involved a variant of the three-alternative forced-choice task used during training on day 1 and 2 (Fig. 1D-E). However, unlike the training task, the background colour switched randomly between trials to indicate either the day 1 or day 2 context, and the task was presented in the absence of feedback. Given the probe stimulus and the background colour, participants were instructed to select the correct associated stimulus. The memory test constituted 100 trials, with half presented on the yellow background and half on the blue background.

### fMRI scan task

The fMRI scan task involved participants viewing the seven visual stimuli used in the training task (1:7), presented via a computer monitor, which for group 1 was then projected onto a screen inside the scanner bore. On each trial two stimuli were presented consecutively for 800 ms each, with an inter-stimulus interval of 300 ms (Fig. 1F). The inter-trial interval was selected from a truncated gamma distribution with mean of 2.9 s, minimum of 1.5 s and maximum of 9.7 s. To control for potential confounding effects of expectation suppression (Summerfield et al., 2008), all stimuli, and all possible pairs of stimuli, were presented equally often in a fully randomised order. Participants were required to perform a task incidental to the contrast of interest which involved identifying whether the presented stimuli were familiar or “oddball.” Oddball stimuli, defined as stimuli that did not belong to the training set 1 to 7, were randomly inserted into *7%* of trials. Participants were instructed to press a button on an MR compatible button box using their right index finger when they identified “oddball” stimuli but not if both stimuli on the trial were familiar. No feedback was given. Each task block included 196 trials and lasted twenty minutes. Each participant performed two task blocks.

### fMRI imaging protocol

Participants in group 1 completed the scan task within a 7 Tesla Magnetom MRI scanner (Siemens) with 1-channel transmit and a 32-channel phased-array head coil (Nova Medical, USA) at the Wellcome Centre for Integrative Neuroimaging Centre (University of Oxford). Current 7T radio-frequency (RF) coil designs suffer from B1 inhomogeneity effects which were pronounced in the right temporal lobe. To overcome this, we positioned two barium titanate dielectric pads (4:1 ratio of BaTiO3:D2O, with a relative permittivity of around ~300, and size 110 × 110 × 5 mm^3^) over the right temporal lobe in all 7T scanning sessions, causing a “hotspot” in the RF distribution at the expense of distal regions (Brink and Webb, 2014; Teeuwisse et al., 2012). The tDCS electrode was situated between the dielectric pad and the head.

To acquire fMRI data a multiband echo planar imaging (EPI) sequence was used to acquire 50 1.5 mm thick transverse slices with 1.5 mm gap, in-plane resolution of 1.5 × 1.5 mm^2^, repetition time (TR) = 1.512 s, echo time (TE) = 20 ms, flip angle = 85°, field of view 192 mm, and acceleration factor of two. The partial volume covered occipital and temporal cortices and in each session 644-723 volumes were collected (~20 min). For each participant, a T1-weighted structural image was acquired to correct for geometric distortions and co-registration between EPIs, consisting of 176 0.7 mm axial slices, in-plane resolution of 0.7 × 0.7 mm^2^, TR = 2.2 s, TE = 2.96 ms, and field of view = 224 mm. A field map with dual echo-time images was also acquired (TE1 = 4.08 ms, TE2 = 5.1 ms, whole-brain coverage, voxel size 2×2×2 mm^3^).

### MRS

For participants in group 1, during the scan session MRS data was acquired as described in (Barron et al., 2016a). B0 shimming was performed in a two-step process. First, GRE-SHIM (field of view, 384 × 384 mm^2^; TR = 600 ms; TE1/2 = 2.04/4.08 ms; slice thickness 4 mm; flip angle 15°; slices 64; scan time 45 s) was used to determine the optimal first-and second-order shim currents. The second step involved only fine adjustment of first-order shims using FASTMAP (Gruetter and Tkác, 2000). The modified semi-LASER sequence, previously shown to have minimal chemical shift displacement error (CSDE), was used with TE = 36 ms, TR = 5’6 s to acquire MRS measurements in a 2 × 2 × 2 cm^3^ volume of interest (VOI), positioned next to the tDCS electrode (Fig. 3D) (Oz and Tkáč, 2011).

For each MRS measurement between 65 and 130 scan averages were collected, giving a total acquisition time of around 10 min. Three measurements were acquired for each participant, before and during tDCS, and after the second task block (Fig. 3A). Metabolites were quantified using LCModel (for example spectra: Supplementary Fig. 3A-B) (Provencher, 1993, 2001). The model spectra of alanine (Ala), aspartate (Asp), ascorbate/vitamin C (Asc), glycerophosphocholine (GPC), phosphocholine (PCho), creatine (Cr), phosphocreatine (PCr), GABA, glucose (Glc), glutamine (Gln), glutamate (Glu), glutathione (GSH), myo-inositol (myo-Ins), Lactate, N-acetylaspartate (NAA), N-acetylaspartylglutamate (NAAG), phosphoethanolamine (PE), scyllo-inositol (scyllo-Ins) and taurine (Tau) were generated based on previously reported chemical shifts and coupling constants by VeSPA Project (Versatile Simulation, Pulses and Analysis) (Govindaraju et al., 2000; Tkac I., 2008).

The unsuppressed water signal acquired from the VOI was used to remove residual eddy current effects and to reconstruct the phased array spectra (Natt et al., 2005). To improve comparability across spectra, the water component of the spectra was then removed before single scan spectra were summed from 32 channels, corrected for frequency and phase variations induced by participants’ motion, and then summed. LCModel analysis was performed on all spectra within the chemical shift range 0.5 to 4.2 ppm (Provencher, 1993).

Reliable LCModel fits were achieved in 20 of the 26 participants and metabolite concentration estimated relative to unsuppressed water spectrum acquired from the same VOI. In the remaining 6 participants GABA quantification was either unreliable or inestimable due to lipid contamination and broader linewidth. The lipid contamination could be observed directly in the spectral range 1.9-0.5 ppm (Supplementary Fig. 3A-B). The broader linewidth, quantified using Full-Width at Half Maximum (FWHM), was significantly higher in these six participants relative to the 20 participants included for analysis (two sample t-test: t_24_=3.33, p=0.001, Supplementary Fig. 3C).

All measured metabolites included in the analysis had Cramér-Rao lower bound (CRLB) ≤ 50% (Bednařík et al., 2015). Relative to baseline concentrations (‘Before tDCS’), the change in GABA (Fig. 3E), glutamate (Fig. 3F), and other metabolite concentrations (Supplementary Table 1) were compared across conditions using a two-tailed paired t-test where the direction of the effect was unknown and a one-tailed paired t-test in instances where the direction of the effect was predicted from previous data (i.e. for the change in GABA).

### tDCS

Immediately before and during block 2 of the scan task, participants in groups 1, 2 and 3 received tDCS using a DC-Stimulator (Eldith) which delivered a 1 mA current to the brain. For group 1, the current was delivered while participants were inside the 7T MRI scanner. For groups 2 and 3, the current was delivered outsider the scanner using a double-blind procedure (see below). To ensure that the tDCS was suitable for use in the 7T scanner, we used two 5 × 7 cm^2^ MRI compatible electrodes (Easycap) fitted with 5 kOhm resistors to minimise the risk of heating or eddy current induction. Using high-chloride EEG electrode gel (Easycap) as a conducting paste, the anodal electrode was placed on the scalp above the region of right temporal cortex previously identified as encoding the association between paired shapes (Fig. 3B-C), approximately at the 10-20 T6 node location. The cathodal electrode was placed over the contralateral supraorbital ridge (Fig. 3B). For participants in group 1, a cod-liver oil capsule was taped to the anodal electrode, immediately underneath the resistor, to make the electrode MR-visible and allow for its location to be mapped onto the anatomical brain surface (Fig. 3D). For all participants, the impedance of tDCS was checked prior to stimulation. In group 1, this impedance check was performed before participants entered the scanner and again once the participant was lying inside the bore of the magnet with extension leads connected to the stimulator. For participants in group 1 and 2, tDCS was delivered using a 10 s ramp-up of the current, which was then held at 1 mA current for a total of 20 min, before a 10s ramp-down. For participants in group 3, sham stimulation involved mimicking the prickling sensation of stimulation using a 10s ramp-up of current, turning stimulation off for 20 minutes and then repeating the 10s ramp-up. For participants in all groups 1-3, the stimulation protocol commenced 10 min prior to the start of the second fMRI scan task (Fig. 3A). At the end of the experiment, participants in groups 2 and 3 were debriefed: they were informed that they may have received sham stimulation and were asked to declare whether they believed they had received real or sham stimulation. The blinded researcher (R.K., see below) also declared whether they believed the participants had received real or sham stimulation.

### Double-blind procedure for delivery of anodal tDCS and sham stimulation

Participants in groups 2 and 3 were first recruited, before being randomly assigned to the anodal (group 2) or sham (group 3) stimulation condition using a random number generator. Randomisation was performed by a researcher (H.B.) who was not involved in recruitment. Behavioural training, electrode placement, the scan tasks, surprise memory test and debrief were carried out by a researcher blind to the stimulation condition (R.K.). tDCS was delivered by a researcher who was aware of the stimulation conditions and who was not involved in any of the behavioural training or assessment (H.B.). Analysis of participants responses during the debrief indicated that 55% of participants in group 2 (‘tDCS’) and 75% of participants in group 3 (‘sham’) believed they received anodal tDCS stimulation. The blinded researcher believed that 55% of participants in group 2 (‘tDCS’) and 45% of participants in group 3 (‘sham’) received anodal tDCS stimulation.

### fMRI data analysis

For all MRI data sets obtained from participants in group 1, pre-processing was carried out using SPM12 (http://www.fil.ion.ucl.ac.uk/spm/). Two participants were excluded from the fMRI analysis due to poor performance on the fMRI scan task (<80% accuracy on one or more of the two task blocks), suggesting that they may have fallen asleep during the task. For the remaining 24 participants images were corrected for signal bias, realigned to the first volume, corrected for distortion using field maps, normalised to a standard EPI template and smoothed using an 8-mm full-width at half maximum Gaussian kernel. For each participant and for each scanning block, fMRI data was analysed in an event-related manner using two different general linear models (GLMs), one designed for univariate analyses and a second one for multivariate analyses. In both GLMs explanatory variables used a delta function to indicate the onset of a trial and were then convolved with the hemodynamic response function.

The first GLM, used to analyse univariate BOLD effects, was applied to data from each of the two scan task blocks separately, and to data from both scan task blocks together. In the design, a total of 46 different explanatory variables were included per block. 42 of these explanatory variables were included to account for each possible pair of visual stimuli (‘1’ and ‘2’, ‘1’ and ‘3’ etc.) in each of the two background contexts (i.e. memory 1 or memory 2), regardless of the order in which the two stimuli were presented within the pair. An additional 4 explanatory variables were used to model trials that included repeating stimuli or trials were ‘odd-ball’ stimuli had been presented in each of the two background contexts (i.e. memory 1 or memory 2). Finally, for each task block an additional 6 scan-to-scan motion parameters produced during realignment were included in the GLM as additional nuisance explanatory variables to account for motion-related artefacts.

Using the output of this first GLM for the univariate analysis, the following three principal contrasts of interest were assessed. First, to measure cross-stimulus adaptation as an index for expression of directly associated stimuli (Fig. 3G-N), the contrast of interest involved comparing fMRI BOLD signal for trials with pairs of stimuli separated by more than one link across both memories (‘unassociated’; i.e. memory 1 and memory 2 links 2-7, 5-7, 2-4, 1-5, 1-4, 2-5, 4-7, 3-6, 1-6, 1-3) with fMRI BOLD signal for trials with pairs of stimuli separated by one link in both memories (‘associated’; i.e. memory 1 and memory 2 links 1-2, 4-5, 7-1). Second, to measure cross-stimulus adaptation as an index for memory interference (Fig. 4A-J), the contrast of interest involved comparing fMRI BOLD signal for pairs of stimuli separated by more than one link across both memories (‘unassociated’; i.e. memory 1 and memory 2 links 2-7, 5-7, 2-4, 1-5, 1-4, 2-5, 4-7, 3-6, 1-6, 1-3) with fMRI BOLD signal for pairs of stimuli separated by more than one link in the current context, but only one link in the alternative context (‘hidden’; i.e. memory 1: links 3-5, 4-6, 2-6, 3-7; memory 2: links 3-4, 5-6, 2-3, 6-7). Third, to measure the BOLD response to trials where there was an opportunity for memory interference (Fig. 2A-C, 2H-I, 4K), the contrast of interest involved comparing fMRI BOLD signal for pairs of stimuli that shared the same topological relationship across the two memories (‘stable’, i.e. links that did not include stimuli 3 or 6) with fMRI BOLD signal for pairs of stimuli that had a different topological relationship across the two memories (‘unstable’; i.e. links that included stimuli 3 or 6).

### ROI specification

To assess fMRI cross-stimulus suppression effects in the neocortex, three ROIs were defined. To assess evidence for replication of previously published results (Fig. 3I-N), an independent ROI was defined using the previously published dataset, after thresholding the contrast of interest at p<0.01 uncorrected (Fig. 3C,3H) (Barron et al., 2016a). To assess evidence for memory interference, an ROI was defined from the peak average location of the anodal tDCS electrode. This was estimated using the T1 scan to identify the location of the cod-liver oil capsule taped immediately underneath the resistor of the anodal electrode. For each participant, the ventral-dorsal coordinate was taken from the upper edge of the cod-liver capsule. The medial-lateral coordinate was projected 20mm from the later surface, consistent with the peak medial-lateral coordinate of previously published cross-stimulus suppression induced by application of tDCS (Fig. 3C) (Barron et al., 2016a). For each participant, an 8mm sphere was then drawn around the identified coordinate, and the sphere warped to a standard EPI template. Across individuals the average ROI was calculated (Fig. 3D). To perform SVC for multiple comparisons, a 10mm sphere was drawn around the peak of the group average tDCS electrode location (Fig. 3M) and cluster-defining threshold set to p<0.001 uncorrected with peak-level FWE corrected at p<0.05. Capitalizing on variance across participants within this larger ROI, the extracted fMRI cross-stimulus suppression measures were correlated with changes in GABA and behaviour.

To assess changes in BOLD signal in the hippocampus (Fig. 2), a hippocampal mask was used with a univariate cluster-defining threshold of p<0.001 uncorrected combined with SVC for multiple comparisons (peak-level FWE corrected at p<0.05).

### fMRI statistics

From the first GLM, the contrast images of all participants were entered into a second-level random effects analysis. To test for fMRI cross-stimulus suppression effects in an unbiased fashion, parameter estimates obtained from the relevant GLM were extracted from an independent region of interest (ROI) (see above). Paired t-tests were used to assess differences in the main effect between sessions. When testing evidence for replication of our previous findings (Barron et al., 2016a) a one-tailed test was used. In all other instances, two-tailed tests were used. The significance level was set to p<0.05.

The second GLM was used to assess multivariate effects. In this GLM each trial was modelled as a unique explanatory variable. In addition, 6 scan-to-scan motion parameters produced during realignment were included in the GLM as additional nuisance explanatory variables to account for motion-related artefacts. The output of this GLM was used to estimate the representational similarity between each trial, using the representational similarity analysis toolbox (RSA) (Kriegeskorte et al., 2008; Nili et al., 2014). The dissimilarity between the response pattern elicited on each trial was estimated using the Mahalanobis distance (Walther et al., 2016), and expressed using correlation distances (1-r). For each participant, the dissimilarity value for the response patterns associated with each trial was represented in each cell of a representational dissimilarity matrix (RDM). Using these RDMs, a confusion matrix was estimated for each participant by sorting trials by stimulus type. Thus, for each stimulus all trials containing the stimulus were included to estimate a stimulus representation, e.g. stimulus 1 in memory 1 included all pairs of stimuli shown on a yellow background that included stimulus 1, i.e. 1-1, 1-2, … 1-7. For each memory this gave seven different representations, which together formed a 14×14 confusion matrix (Fig. 2I). This confusion matrices was used to estimate summary statistics, which were tested at the group level using a two-sided signed-rank test across participants (Wilcoxon 1945). This indicated whether the difference in correlation coefficients between two conditions was greater than zero.

### Correlations with fMRI data

To assess the relationship between hippocampal BOLD signal and behaviour, a Pearson’s correlation was used. Due to outlier data points in the fMRI cross-stimulus suppression measure (see Fig. 3N, Fig. 4B-D, Fig. 4I-J), Spearman’s rank correlation was used to assess the relationship between fMRI cross-stimulus suppression and changes in GABA or behaviour. Correlations were plotted between standardized residuals, using a partial correlation method to control for the effect of other variables. To aid interpretability, partial correlation plots in Fig. 2-4 show the mean of the original variables added to the standardized residuals.

A permutation test was used to quantify the difference in correlation between behavioural performance and hippocampal BOLD in block 1 versus block 2. To estimate a null distribution subject labels for hippocampal BOLD signal were permuted 10,000 times, before being correlated with behavioural performance. The difference in correlation between the block 1 and block 2 correlations was then computed for all 10,000 examples. The true difference between block 1 and block 2 correlations was compared against the null distribution to generate a p-value (Supplementary Fig. 4D).

### Data and code availability

Upon publication Matlab scripts for reproducing all figures will be made available on GitHub (https://github.com/rskool/meminf). Upon publication group t-stat images, anonymized subject-specific parameter estimates extracted from ROIs, and relevant experimental parameters that support the findings of this study will be made available on GitHub (https://github.com/rskool/meminf).

